# European and tropical *Aedes albopictus* mosquito populations have similar systemic Zika virus infection dynamics

**DOI:** 10.1101/764498

**Authors:** Sébastian Lequime, Jean-Sébastien Dehecq, Sébastien Briolant, Albin Fontaine

## Abstract

First isolated from a forest in East Africa in the mid-20^th^ century, Zika virus (ZIKV) has now emerged worldwide in urbanized areas where its mosquito vectors, mainly *Aedes aegypti* and *Ae. albopictus*, are present. Europe and French overseas territories in the Indian Ocean have been so far spared despite the presence of *Ae. albopictus*, the Asian tiger mosquito. However, because they have strong economic and touristic links with regions affected by ZIKV, French territories in the Indian Ocean have a high risk of introduction. Here, we assess the susceptibility of two *Ae. albopictus* populations from Metropolitan France and the Reunion island (a French oversea territory in the Indian Ocean) for a ZIKV isolate from the Asian genotype at a titer ranging from 3 to 7.5 × 10^6^ focus-forming units per milliliter. High infection rates and unpreceded levels of systemic infection rates were observed in both Metropolitan France and the Reunion island populations, without differences in infection rates or intra-mosquito systemic infection dynamics between the two mosquito populations. Ten and 20-days were needed by the virus to disseminate in 50% and 100% of the exposed mosquitoes respectively. Such slow intra-mosquito viral dynamics, in addition to repeatedly reported high transmission barrier in the literature, can impact ZIKV transmission when potentially vectored by *Ae. albopictus*. However, because mosquito-borne virus intra-host transmission dynamics can be influenced by numerous factors, including virus dose dynamics inside infectious humans or viral evolution towards shorter extrinsic incubation periods (EIP), our results highlight that *Ae. albopictus* populations present in Metropolitan France and the French territoires in the Indian Ocean might become potential vector for autochthonous ZIKV transmissions.

## Introduction

Several mosquito-borne viruses have left their original enzootic cycles in tropical primary forests to emerge worldwide in transmission cycles involving humans and mosquitoes highly adapted to urban environments. While *Aedes aegypti* is considered as the main vector species of viruses affecting human health, the Asian tiger mosquito *Aedes albopictus* is trailing second in the list and might soon move ahead because of its outstanding invasive capacity. Originating from South-East Asia, *Ae. albopictus* has invaded the world and is now present in all continents, including temperate Europe, due to its potential of enduring harsh winter conditions through the induction of diapause in eggs^1,2^. The species is also displacing *Ae. aegypti* populations in areas where both species co-localize due to competitive advantages, notably at the larval stage^3–5^.

Zika virus (ZIKV) is the lastest mosquito-borne virus that has emerged on a global scale and that can be sustained in urban cycles involving solely human-mosquito transmissions. ZIKV is a RNA virus from the *Flavivirus* genus, Flaviviridae family, which includes other human pathogens such as yellow fever virus (YFV) or dengue viruses (DENV). First isolated from a rhesus monkey in the Zika forest in Uganda in 1947, ZIKV has emerged concomitantly in 2007 in Gabon^6^ and in the Federated States of Micronesia where it triggered a major outbreak in Yap Island^7^. The virus then spread through the South Pacific islands from 2013 to 2014^8–10^ before its emergence in north-eastern Brazil in 2015^11^, the starting point toward a large outbreak that hit a total of 50 territories and countries in the Americas^12^. Usually relatively mild, ZIKV infection can impact human health by leading to Guillain-Barré syndromes and cases of microcephaly in newborns, as well as other neurological impairments^13^.

*Aedes albopictus* mosquitoes are known to vector several major arthropod-borne viruses (arboviruses)^14^ and have been incriminated in the transmission of chikungunya (CHIKV) and DENV viruses in major recent outbreaks in the Reunion island, a French oversea territory in the Indian Ocean^15,16^. This species was also responsible for the transmission of these viruses in Europe (e.g. autochthonous cases in South France^17–19^) and a CHIKV outbreak in north-eastern Italy with over 200 confirmed cases^20^. European French populations of *Ae. albopictus* mosquitoes were shown to experimentally transmit DENV and CHIKV as efficiently as the typical tropical vector *Ae. aegypti*^21^. On the other hand, ZIKV has exhibited lower dissemination and transmission rates in experimental exposure assays in both *Ae. albopictus* and *Ae. aegypti* when compared to other RNA mosquito-borne viruses such as DENV, YFV or CHIKV ^22–24^. Such barriers preventing systemic infection (virus dissemination from midgut to secondary tissues) or transmission^22,24–27^ suggest limited risk of Zika virus transmission in Europe or the Reunion island^22,25^. To our knowledge, no ZIKV outbreak or autochthonous transmission was yet reported in Metropolitan France or French overseas territories where this species is established, but the implication of *Ae. albopictus* was strongly suspected in West Africa^6^. French overseas territories in the Indian Ocean have repeatedly been hit by mosquito-borne viruses and a large amount of goods and travelers are transiting every year between Metropolitan France and these tropical overseas territories, putting these regions at risk of ZIKV emergence.

In the present studies, we aim to assess the susceptibility of *Aedes albopictus* populations from Metropolitan France and Oversea France (the Reunion island, Indian Ocean) for ZIKV infection and compare their intra-host systemic infection dynamics. Two *Ae. albopictus* mosquito populations were orally exposed to a 3 to 7.5 ×10^6^ focus-forming units per mL (FFU/mL) infectious blood meal of a ZIKV isolate from the Asian genotype. These virus titers lie between median infection doses that were reported for *Ae. albopictus*^27^. Mosquitoes from each population were sampled at five different times post virus exposure to assess intra-mosquito systemic infection dynamics. This assessement of intra-mosquito systemic infection dynamics of ZIKV for both tropical and temperate *Ae. albopictus* mosquito populations shed light on the risk of ZIKV emergence in both temperate and tropical areas.

## Materials and Methods

### Zika virus isolate

In this study, we used the ZIKV isolate SL1602 that was isolated from the plasma of a traveller returning from Suriname in 2016^28^. The full-length consensus genome sequence of the isolate is available from GenBank under accession number KY348640. This isolate belongs to the Asian lineage and is phylogenetically close to other viruses that have recently been isolated in the Americas^28^. The virus was passaged four times in C6/36 cells (*Aedes albopictus*) prior use for mosquito infections. To prepare virus stock, sub-confluent C6/36 cells were infected using a viral multiplicity of infection (MOI) of 0.1 (one infectious viral particle for 10 cells) in 25-cm^2^ culture flasks and incubated for 7 days at 28 °C with 5 mL of Leibovitz’s L-15 medium supplemented with 0.1% penicillin (10,000 U/mL)/streptomycin (10,000 μg/mL) (Life Technologies, Grand Island, NY, USA), 1X non-essential amino acids (Life Technologies) and 2% fetal bovine serum (FBS, Life Technologies). At the end of the incubation, the cell culture medium was harvested, adjusted to 10% FBS and pH ∼8 with sodium bicarbonate, aliquoted and stored at −80 °C. An additional 5 mL of Leibovitz’s L-15 medium prepared as described above were added to infected confluent C6/36 cells and harvested 3 days later. This procedure increases virus titres in the stock solution.

Virus titration was performed by focus-forming assay (FFA) on one aliquot that has been stored at −80 °C as previously described with minor modifications^29^. This assay relies on inoculation of 10-fold dilutions of a sample onto a sub-confluent culture of C6/36 cells, followed by incubation and subsequent visualization of infectious foci by indirect immunofluorescence. Modifications to the previously published protocol include an 1 hour incubation step at 37 °C with 40 μL/well of mouse anti-*Flavivirus* group antigen antibody clone D1-4G2-4-15 (Merck Millipore, Molsheim, France) diluted 1:200 in PBS + 1% bovine serum albumin (BSA) (Interchim, Montluçon, France). After another three washes in PBS, cells were incubated at 37 °C for 30 min with 40 μL/well of a goat anti-mouse IgG (H+L), FITC-conjugated antibody (Merck Millipore). After three more washes in PBS and a final wash in ultrapure water, infectious foci were counted under a fluorescent microscope and converted into focus-forming units/mL (FFU/mL). The titre of the frozen virus stock was estimated at 6.35 × 10^5^ focus-forming units per mL (FFU/mL) for the first medium culture harvest and 2.75 × 10^8^ FFU/mL for the second medium culture harvest. All infectious experiments were conducted in a Biosafety Level-3 (BSL-3) insectary (IHU Méditerranée Infection, Marseille).

### Mosquito populations

Two mosquito populations of *Aedes albopictus* species were used in this study, one from Metropolitan France and the other from the Réunion island, a French overseas territory in the Indian Ocean. *Ae. albopictus* mosquitoes from Metropolitan France were collected both as eggs using ovitraps and human landing catches in three different locations in Marseille city (5^th^, 12^th^ and 13^th^ districts) in June 2018. *Ae. albopictus* mosquitoes from French overseas territory were all collected as eggs in June 2018 from 6 different localities of the Réunion island named Sainte Marie Duparc, Saint André centre-ville, Saint Paul, Le Port Rivière des Galets, Saint Leu Pointe des châteaux, Saint Pierre Bois d’olive (Figure 1). An first generation population was obtained from the eggs of adults originating from ovitraps or captured by human landing catches. For each population, larvae from different collection sites were combined and adult mosquitoes were maintained in standardized insectary condition (28°C, 75 ± 5% relative humidity, 16:8 hours light-dark cycle) until the 4^th^ generation by mass sib-mating and collective oviposition. Adult females from each population were blood-fed on human blood obtained from the Etablissement Français du Sang (EFS) through a membrane feeding system (Hemotek Ltd, Blackburn, UK) using pig intestine as membrane. Access to human blood was based upon an agreement with the EFS. Eggs were hatched in reverse osmosis water and larvae were reared with a standardized diet of fish food in 24 × 34 × 9 cm plastic trays at a density of about 400 larvae per tray. A maximum of 800 male and female adults were maintained in 24 × 24 × 24 cm screened cages with permanent access to a 10% sucrose solution.

**Figure 1:**
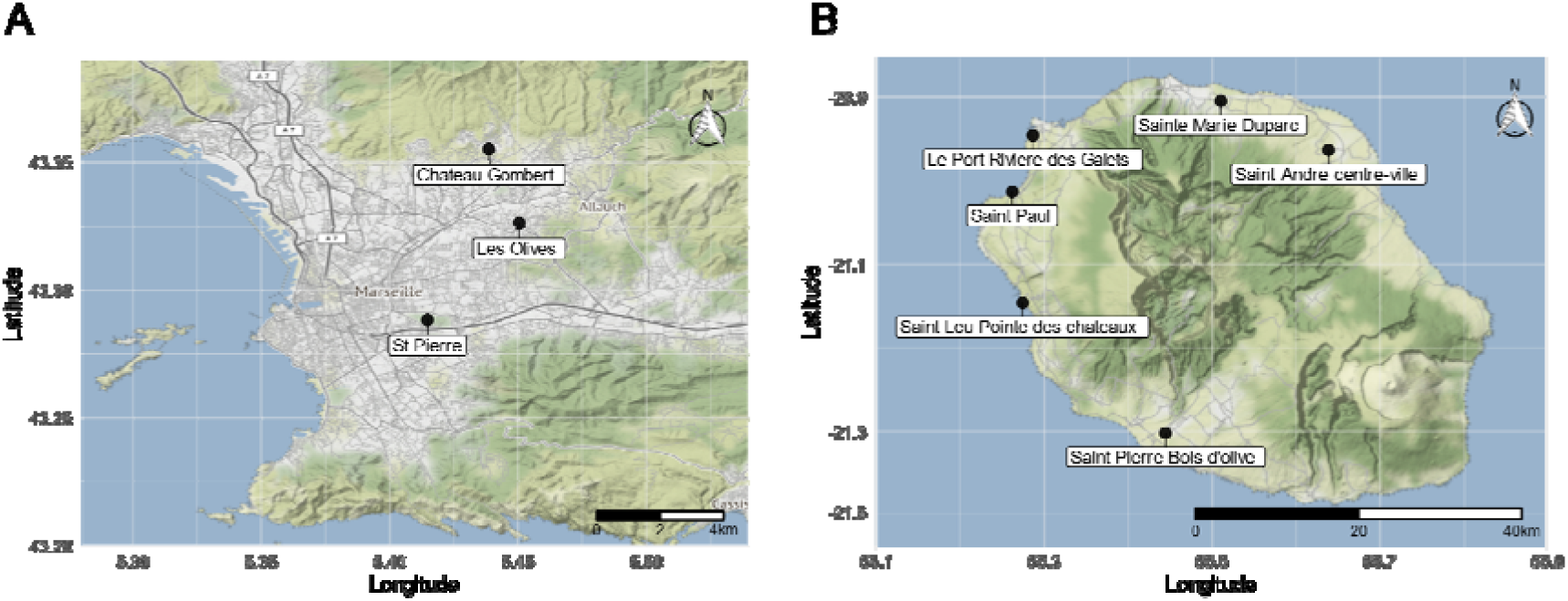
Geographic location of the F0 mosquito collecting sites in Marseille (A) and Reunion island (B). Mosquitoes from these sites were used as progenitors of the Metropolitan France and Overseas France populations (4^th^ generation) that were used in this study.

### Experimental mosquito infections

Nine to 13 days old females were transferred to the biosafety level 3 laboratory and deprived of sucrose solution 24 hours before experimental virus exposure. Sixty to 80 females were confined into 80-mm high and 80/84 mm in (inside/outside) diameter cardboard containers. Containers were sealed on the top with mosquito mesh and with a 65-mm high polystyrene piston covered up with a plasticized fabric that match the inside diameter of the container at the bottom. Pistons of each container were rose before to infectious blood meal exposure to contain all mosquitoes in a tight space below the infectious blood meal. This procedure increases the yield of engorged mosquitoes. Females were allowed to feed for 20 min from an artificial feeding system (Hemotek) covered by a pig intestine membrane that contained the infectious blood meal maintained at 37 °C. Feeders were placed on top of the mesh that sealed the containers. The infectious blood meal consisted of two volumes of washed human erythrocytes and one volume of viral suspension. Human erythrocytes were collected and washed one day before the experimental infection. An aliquot of the artificial blood meal was collected immediately prior to blood feeding and titred by FFA as described above without a freezing step. Final ZIKV virus titres in blood meals were 7.5 × 10^6^ FFU/mL and 3 × 10^6^ FFU/mL for experiment 1 and 2, respectively. After virus exposure, fully engorged females were cold anesthetized and sorted on ice before being individually transferred to new cardboard containers and maintained under controlled conditions (28°C, 75 ± 5% relative humidity, 16:8 hours light-dark cycle with permanent access to a 10% sucrose solution).

### ZIKV RNA detection

ZIKV RNA was detected in mosquito bodies and heads after a fresh dissection from freeze-killed mosquitoes at 5,10, 14, 17 and 21-days post virus exposure (DPE) for both *Ae. albopictus* populations. Presence of ZIKV RNA in bodies indicates a midgut infection, while presence of the virus RNA in mosquito heads indicates a systemic (disseminated) infection^30^. These two vector competence indices were determined qualitatively (i.e. presence or absence of virus in mosquito bodies and heads, respectively). Dissected mosquito heads and bodies were homogenized individually in 400 μL of lysis buffer (NucleoSpin® 96 Virus Core Kit, Macherey-Nagel, Hoerdt, France) during three rounds of 20 sec at 5,000 rpm in a mixer mill (Precellys 24, Bertin Technologies, Montigny le Bretonneux, France). Viral RNA from individual organs was extracted using the NucleoSpin® 96 Virus Core Kit (Macherey-Nagel) according to the manufacturer’s instructions. At the final step, viral RNA from each sample was eluted in 100 μL of RNase-free elution buffer.

Detection of ZIKV RNA was performed with an end-point reverse transcription polymerase chain reaction (RT-PCR) assay. ZIKV genomic RNA was first reverse transcribed to complementary DNA (cDNA) with random hexamers using M-MLV Reverse Transcriptase (Life Technologies) according to the manufacturer’s instructions. cDNA was amplified by 35 cycles of PCR using the set of primers targeting the RNA-dependent RNA polymerase NS5 gene: ZIKV-F: 5’-GCCATCTGGTATATGTGG-3’ and ZIKV-R: 5’-CAAGACCAAAGGGGGAGCGGA-3’. Amplicons (393 bp) were visualized by electrophoresis on 1.5% agarose gels.

### Statistical analysis

The time-dependent effect of the mosquito population on mosquito body infection and systemic infection was analysed by Firth’s penalized likelihood logistic regression by considering each phenotype as a binary response variable. Penalized logistic regression, implemented through the *logistf* R package^31^ was used to solve problem of separation that can occur in logistic regression when all observations have the same event status for a combination of predictors or when a continuous covariate predict the outcome too perfectly. A full-factorial generalized linear model that included the time post-virus exposure and the mosquito population was fitted to the data with a binomial error structure and a logit link function. Statistical significance of the effects was assessed by an analysis of deviance.

The intra-host dynamic of systemic infection was assessed by a global likelihood function for each mosquito species as described by Fontaine *et al*, 2018^30^. Probabilities of systemic infection at each time point post virus exposure were estimated with a 3-parameter logistic model. The probability of systemic infection at a given time point (*t*) is governed by *K*: the saturation level (the maximum proportion of mosquitoes with a systemic infection), *B*: the slope factor (the maximum value of the slope during the exponential phase of the cumulative function, scaled by *K*) and *M*: the lag time (the time at which the absolute increase in cumulative proportion is maximal). For easier biological interpretation, *B* was transformed into a rising time Δt, which correspond to the time required to rise from 10% to 90% of the saturation level with the formula: Δt = ln (81) / *B*. For each mosquito population, the *subplex*^32^ R function was used to provide the best estimates of the three parameters to maximize the global likelihood function (i.e., the sum of binomial probabilities at each time point post virus exposure). This method accounts for differences in sample size when estimating parameters values.

All statistical analyses were performed in the statistical environment R v3.5.2^33^. Figures were prepared using the package ggplot2^34^.

## Results

A total of 227 and 175 engorged female mosquitoes from the Metropolitan France and Overseas France populations, respectively, were analyzed in this study. Two ZIKV experimental exposure assays were performed with infectious blood meals titred at 7.5 × 10^6^ FFU/mL (98 and 120 engorged mosquitoes for Metropolitan France and Overseas France populations, respectively) and 3 × 10^6^ FFU/mL (129 and 55 engorged mosquitoes for Metropolitan France and Overseas France populations, respectively). Because each assay was performed with a single dose, the virus dose effect was here confounded with the experiment effect. High body infection prevalences were obtained for both populations, independently of the time post exposure (Figure 2). Averaged over time, mean body infection prevalences were 84% for the Metropolitan France population for each experiment and over 87% for the Overseas France population. A full factorial logistic regression model revealed no statistically significant difference for body infection prevalences across mosquito populations and experiment replicates. However, there was a statistically significant interaction effect between time post-virus exposure and the experiment replicate (analysis of deviance, p-value = 0.048) which was driven by a relatively low body prevalence at 5 days post-virus exposure for mosquitoes from the Metropolitan France population at 7.5 × 10^6^ FFU/mL.

**Figure 2:**
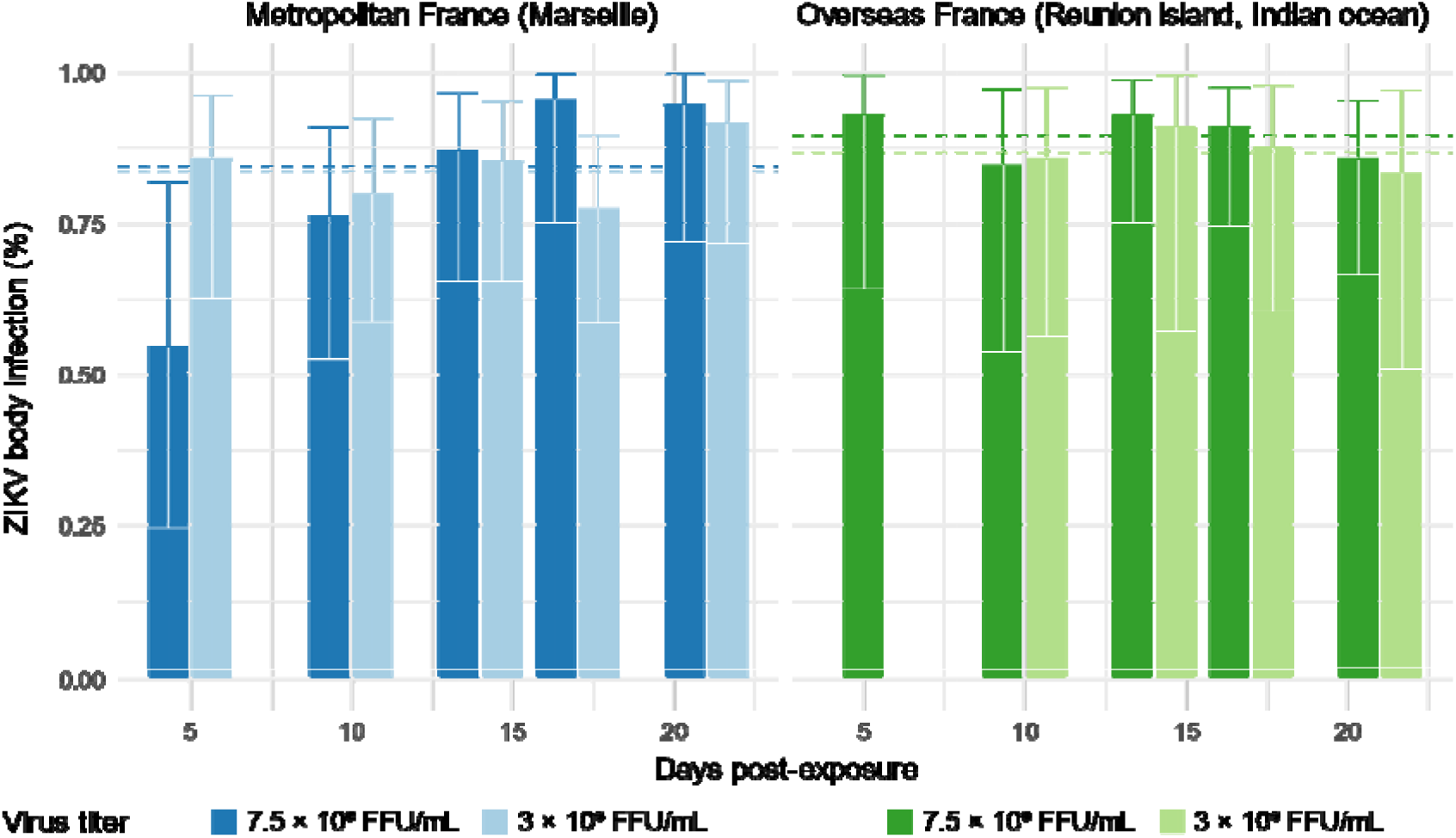
Body infection prevalence for *Ae. albopictus* mosquitoes from two populations exposed the same isolate of ZIKV at two different titers. Percentages of body infections over time post-ZIKV exposure are represented for the Metropolitan France population (left panel, blue color) and Overseas France population (right panel, green color) with their 95% confidence intervals. ZIKV titers at which each mosquito population was exposed to are represented with a dark (7.5 × 10^6^ FFU/mL) and light (3 × 10^6^ FFU/mL) hue in each panel.

**Figure 3:**
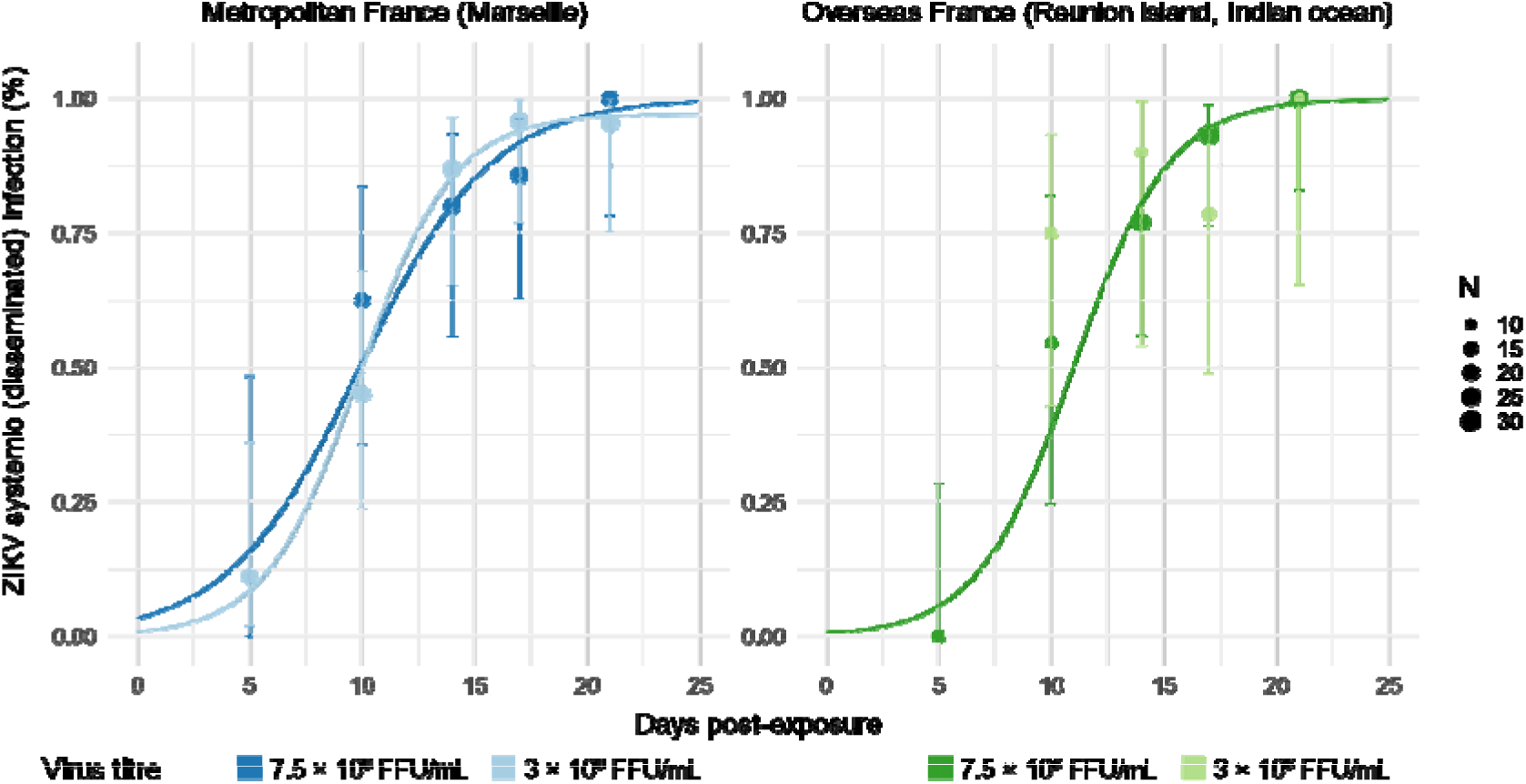
Systemic infection kinetic for *Ae. albopictus* mosquitoes from two populations exposed the same isolate of ZIKV at two different titers. Cumulative prevalences of systemic (disseminated) infections over time post ZIKV exposure are represented as points for the Metropolitan France population (left panel, blue color) and Overseas France population (right panel, green color). Dashes represent the 95% confidence intervals of prevalences. Points size indicates the number of samples (N). The fitted values obtained with a 3-parameter logistic model are represented for each population by a line. ZIKV titers at which each mosquito population was exposed to are represented with a dark (7.5 × 10^6^ FFU/mL) and light (3 × 10^6^ FFU/mL) hue in each panel. No line is represented for the Overseas France population at 3 × 10^6^ FFU/mL due to a model failure for this condition that was caused by a lack of samples at early times post-virus exposure.

Systemic infection prevalences were measured by counting the number of mosquitoes with infected heads over the number of mosquitoes with an infected body. Systemic infection prevalences were significantly influenced by the time post-exposure (analysis of deviance, p-value < 2 ×10^−16^) but not by mosquito population or experiment replicate. Intra-host dynamic of systemic ZIKV infection was inferred in all combinations of mosquito population and virus doses by fitting a 3-parameters logistic model that assumes a sigmoidal distribution of the cumulative proportion of mosquitoes with a systemic infection over time. The model failed to provide relevant estimates for the Overseas France population at the lowest virus dose due to a lack of samples at early times post-virus exposure. However, systemic infection prevalences saturated at 100% for both mosquito populations and experiment replicates. In addition, the model inferred a lag time (M, the inflexion point of the sigmoid which represents the time needed to reach 50% of the saturation level) and a rising time (Δt, time to go from 10% to 90% of the saturation level). The estimated time required to reach 50% of systemic infections was 10 and 11 days for the Metropolitan France and the Overseas France populations, respectively, independently of the experiment replicate for the Metropolitan France population. In the first experiment replicate (highest virus titer at 7.5 × 10^6^ FFU/mL), an estimated 9 and 13 days were needed to go from 10% to 90% of systemic infections for the Overseas France and Metropolitan France populations, respectively. This rising time (Δt) was estimated to 9 days for the Metropolitan France population in the second experiment replicate (virus titer at 3 × 10^6^ FFU/mL).

## Discussion

The French overseas territories in the Indian Ocean have housed several major mosquito-borne virus outbreaks over the last fifteen years. From 2005 to 2006, chikungunya virus has emerged in the Indian Ocean, causing 244,000 cases in the Reunion island, which represented 40% of the island population. Fifteen years later, this region was affected by a dengue virus outbreak that caused more than 6,500 cumulated cases from austral summer 2017 to winter 2018^16^. Neighbouring islands, such as the Seychelles, Mayotte and Mauritius, were also affected^16,35^. These regions are popular destinations for tourists, especially those coming from Metropolitan France. In 2018, the Reunion island hosted more than 400,000 visitors from Metropolitan France, which represents around 80% of the island global tourism flow of the year^36^. This region also has historic, touristic and economic links with countries in the Indian Ocean basin, including India and south-eastern African countries. Chikungunya viruses that were responsible for the 2005-2006 outbreak in the Reunion island were tracked back to Kenya^37^ and then propagated to India^38^. The DENV epidemic that occurred in both the Reunion island and the Seychelles in 2018 was suspected to involve virus strains originating from India or China^16^. To our knowledge, no Zika virus outbreak has been documented in the Reunion island or surrounding islands. Yet, Zika viruses from the Asian lineage were circulating in India from 2016 to 2018^39^, transmitted by the mosquito *Ae. aegypti*^40^. The risk of ZIKV introduction in the Reunion island and surrounding islands is therefore significant.

ZIKV was isolated from *Aedes africanus* mosquitoes in 1948 in the Zika Forest, Uganda, one year after its first isolation from a sentinel rhesus monkey in the same area^41^. While *Aedes hensilli* was strongly suspected to transmit ZIKV during the Yap island outbreak^7^, the species *Aedes aegypti* was incriminated in all major outbreaks that have occurred in the Pacific Ocean, Brazil or the Caribbean, especially in urban transmissions^42^. *Ae. albopictus* was found to be competent for ZIKV experimentally^24,27,43–46^ and strongly suspected to sustain a ZIKV epidemic activity (African lineage) in Gabon in 2007^6^. *Ae. albopictus* has been implicated in all arbovirus transmissions occurring in the Reunion island and could thus vector ZIKV after an introduction in the island. Southern Europe is also at risk in areas where *Ae. albopictus* is implanted, which represents 42 (43.75%) over 96 Metropolitan French departments in 2018^47^. To our knowledge, no autochthonous cases of ZIKV has occurred in Metropolitan France despite the presence of imported cases^48^, while several cases of autochthonous CHIKV and DENV were repeatedly reported during the last fifteen years^17,18,49^.

Here we described high infection rates and unpreceded levels of systemic infection rates that reach 100% in both Metropolitan France and Indian Ocean populations. Importantly, we considered the dynamic nature of vector competence by assessing systemic infection (dissemination) rate over time post-virus exposure. No differences in infection rates or intra-mosquito systemic infection dynamics were revealed between *Ae. albopictus* populations originating from Metropolitan France and the Indian Ocean (the Reunion island). Vector competence studies performed on *Ae. albopictus* often revealed a low systemic infection rate for ZIKV^24,25,43,44,50^. All these studies used a ZIKV isolate from the Asian genotype and a virus dose in the same range than our study (from 3 × 10^6^ FFU/mL to 10^7^ FFU/mL; or 10^7^ TCID_50_/mL) and some have used mosquito populations that can be assumed to be closely related to our *Ae. albopictus* populations (Indian Ocean or European origins). In addition, three of these works assessed systemic infection prevalence at a time that exceed our estimated saturation level (*i*.*e*. 21 days post virus exposure). Mosquito vector competence for viruses were reported to differ according to mosquito populations^51,52^, virus isolates^30^, their combinations^53^ or the virus dose^27^. More complex interactions between intrinsic (*eg*. mosquito virome^54^, endogenous non-retroviral elements^55^ or bacterial microbiome^56^) or environmental (*eg*. temperature^57–59^, rearing or experimental settings) conditions might further complicate comparisons across vector competence studies. An almost unanimous consensus is however prevailing on the existence of a low transmission efficiency^22,24,25,27,43,44,50,60^ in the *Ae. albopictus*/ZIKV couple. Forced salivation assays allow to determine viable infectious particles in single mosquito saliva, avoiding the use of animal experiments. In these assays, mosquito wings and legs are removed prior to collect the saliva from the remaining portion of live mosquito by inserting the proboscis into a capillary filled with different types of media, including defibrinated blood, fetal bovine serum or mineral oil^61,62^. Transmission dynamic estimates must be considered with caution because the method to detect viruses in mosquito saliva might not distinguish true negatives from negatives resulting from mosquito that did not expectorate saliva. This can be however improved by the visualization of the saliva droplet with the use of a hydrophobic medium. Further experiments are needed to assess the presence of a transmission barrier in our settings. It is worth noting that on the contrary, two studies have revealed high transmission rates of ZIKV in *Ae. albopictus*^45,46^.

Interestingly, our results show that ZIKV has slower intra-host systemic infection dynamics than other mosquito-borne viruses: by comparison, 100% systemic infections of *Aedes aegypti* can be observed for several dengue isolates around 10 days (2 folds faster)^30^. Combined with vector longevity, intra-host mosquito dynamic is the most powerful contributor to vectorial capacity, a restatement of the basic reproductive rate (R_0_) of a pathogen ^30,63^. In presence of a ZIKV outbreak sustained solely by *Ae. albopictus* mosquitoes, older individuals (>20 days post their first infectious blood meal) would ultimately be implied in virus transmission and any anti-vector measures designed to reduce mosquito life span should thus have significant effect to control the outbreak.

The efficient infection of *Ae. albopictus* however leaves the opportunity for ZIKV to evolve towards shorter intra-mosquito dynamics, as had already occurred in other systems, enhancing the epidemic potential of the *Ae. albopictus*/ZIKV couple^64,65^. In addition, intra-mosquito systemic infection dynamics and the duration of the extrinsic incubation periods (EIP) depend on the amount of virus ingested during the blood meal^29,66^. ZIKV viremia was shown to peak during the first days of infection and to decline quickly in the next few days when assessed on nonhuman primate models^67,68^. In humans, the amount of ZIKV genomic RNA was only assessed post-symptom onsets: ZIKV genomic RNA copies per millilitre were at their highest level at the first sampling time post-symptom onsets but rarely exceeded 6 logs^69–71^, which would translate into a much lower concentration of infectious particles due to a genomic RNA/infectious particles ratio > 1^72^. At these dose levels the probability to infect *Ae. albopictus* mosquitoes would be minimal, and long EIP values distribution would probably arise. We currently lack information about the distribution of infectious ZIKV concentration in human before symptom onsets. Here, we estimated that around 10 days are required to reach 50% of systemic infections in *Ae. albopictus* at a dose ranging from 3 to 7.5 × 10^6^ FFU/mL. We can hypothesize that faster intra-mosquito virus dynamic could be achieved at a higher virus doses and that infected humans would better contribute to ZIKV transmission in their early days’ pre-symptom onsets. Further works are needed to assess the ZIKV dose impact on intra-mosquito systemic and transmission dynamics.

Taken together, our results suggests that despite being reported as a relatively poor vector of ZIKV, *Ae. albopictus* mosquitoes present in Metropolitan France and French territories in the Indian Ocean basin cannot be considered *per se* as an obstacle to autochthonous ZIKV transmissions. Further works on the assessment of a transmission barrier for ZIKV and the influence of the virus dose on intra-host dynamics are clearly needed to explore the full epidemic potential of the ZIKV/*Ae. albopictus* couple.

## Funding statement and acknowledgments

This study was funded by the Direction Générale de l’Armement (grant no PDH-2-NRBC-2-B-2113) and supported by the European Virus Archive goes Global (EVAg) project that has received funding from the European Union’s Horizon 2020 research and innovation program under grant agreement N° 653316. The contents of this publication are the sole responsibility of the authors. The funders had no role in study design, data collection and interpretation, or the decision to submit the work for publication. Sebastian Lequime is funded by a postdoctoral grant of the Fonds Wetenschappelijk Onderzoek – Vlaanderen (FWO). We thank Nathalie Wurtz, Muriel Militello and Jean-Marc Feuerstein for their help and contributions concerning the development of the biological safety level 3 procedure settings concerning experimental mosquito exposure to ZIKV. We also thank Joël Mosnier and Isabelle Fonta for their help with the seeting up of BSL3 experiments.

## Conflict of interests

The authors declare that there is no conflict of interest regarding the publication of this article.

